# Gene co-expression networks identify novel candidate genes for moulting and development in the Atlantic salmon louse (Lepeophtheirus salmonis)

**DOI:** 10.1101/2021.04.23.441111

**Authors:** Zhaoran Zhou, Christiane Eichner, Frank Nilsen, Inge Jonassen, Michael Dondrup

## Abstract

**Background:** The salmon louse (*Lepeophtheirus salmonis*) is an obligate ectoparasitic copepod, living on Atlantic salmon and other salmonids in the marine environment. Salmon lice cause a number of environmental problems and lead to large economical losses in aquaculture every year. In order to develop novel parasite control strategies, a better understanding of the mechanisms of moulting and development of the salmon louse at the transcriptional level is required.

**Methods:** Three weighted gene co-expression networks were constructed based on the pairwise correlations of salmon louse gene expression profiles at different life stages. Network-based approaches and gene annotation information were applied to identify genes that might be important for the moulting and development of the salmon louse. RNA interference was performed for validation. Regulatory impact factors were calculated for all the transcription factor genes by examining the changes in co-expression patterns between transcription factor genes and deferentially expressed genes in middle stages and moulting stages.

**Results:** Eight gene modules were predicted as important, and 10 genes from six of the eight modules have been found to show observable phenotypes in RNA interference experiments. We knocked down five hub genes from three modules and observed phenotypic consequences in all experiments. In the infection trial, no copepodids with the RAB1A-like gene knocked down were found on fish, while control samples developed to chalimus-1 larvae. Also, a FOXO-like gene obtained highest scores in the regulatory impact factor calculation.

**Conclusions:** We propose a gene co-expression network-based approach to identify genes playing an important role in the moulting and development of salmon louse. The RNA interference experiments confirmed the effectiveness of our approach and demonstrated the indispensable role of RAB1A-like gene in the development of salmon louse. In addition to salmon louse, this approach could be generalized to identify important genes associated with a phenotype of interest in other organisms.

## Background

Copepods have been suggested as the most abundant multicellular animal group, with important roles in marine ecosystems [1, 2]. The salmon louse (*Lepeophtheirus salmonis*) is an ectoparasitic copepod on salmonids, with a life cycle that has eight developmental stages separated by moulting, consisting of two nauplius stages, one copepodid stage, two chalimus stages, two preadult stages and the adult stage [3, 4]. Salmon lice are a major challenge to cage-based aquaculture of salmonids and cause large economical losses each year [5]. The emergence of salmon lice resistances against several drugs makes the situation even worse [6, 7]. Developing novel antiparasitic strategies is thus an urgent and vital issue. To achieve this, we require a thorough understanding of the molecular mechanism of life stages development of the salmon louse. Identifying key genes that influence or regulate the lifespan in the salmon louse is also of great importance for detecting novel drug targets against salmon lice.

Moulting, the shedding and replacement of the exoskeleton common to arthropods, plays a crucial role in the survival and development of copepods. Moulting consists of different events, including detachment of the old cuticle, synthesis of new cuticle, shedding of the old cuticle, hardening of the new cuticle and absorption of the old cuticle. In the salmon louse, moulting occurs cyclicly during development until the adult stage [8, 9]. Steroid hormones such as 20-hydroxyecdysone (20E) play a crucial role in arthropod moulting by regulating a series of pathways [10, 11]. The binding of 20E to ecdysone receptors (EcR) leads to cascade controlling of moulting. The polysaccharide chitin and other structural proteins are the major components of the exoskeleton, so genes involved in the chitin metabolism are supposed to be important for moulting. Particular attention had been devoted to the mechanism of copepods moulting at the molecular level [12, 13], and there have been some studies on the role of steroid hormones and ecdysone receptors in the moulting of salmon louse [14, 15]. Recent investigations on the impact of chitin synthesis inhibitors, compounds belonging to the benzoylurea family (namely diflubenzuron, lufenoron, teflubenzuron, etc.), demonstrate the importance of chitin metabolism for parasite survival and as a target for pest management [16, 17]. Still, there is only limited and often circumstantial knowledge on the molecular mechanisms driving developmental processes in copepods.

In recent years, high-throughput technologies have enabled us to study a large number of genes in parallel and thus facilitate the study of complex biological systems [18]. Being tremendously successful, high-throughput sequencing produces large volumes of data and has enabled a new era of genome research [19]. Our group has recently performed a comprehensive transcriptome time-series analysis using RNA sequencing data from three developmental stages of salmon lice (chalimus-1, chalimus-2 and preadult-1) [8] wherein we applied a method for improved developmental staging of samples by instar-age [20]. That way, we identified genes that may regulate development in this parasite.

A research area that is particularly important for systems biology is the study of dynamic interfaces and crosslinks between different processes and components of biological systems [21]. Recently, a great deal of attention has been devoted to the area of network-based analysis. Network analysis provides a powerful framework for studying a large number of interactions among biological processes and components. Gene co-expression networks (GCNs) have been widely used to capture and mine the interactions among components of the transcriptome [21, 22].

Signatures of hierarchical modularity have been suggested to be present in all cellular networks investigated so far, ranging from metabolic to protein–protein interaction and regulatory networks [23]. In gene co-expression networks, modules are defined as groups of genes with similar expression patterns and can be identified by using clustering methods [24–26]. GCN modules have facilitated a better understanding of a number of biological phenomena [24, 27, 28], and an increasing number of studies based on GCN have been conducted to identify condition-specific gene modules and predict potential genes involved in a certain phenotype [29–32]. In this study, by re-analyzing the staged time-series data produced by Eichner *et al*. [8], we aim at providing a framework for identifying important genes through GCN analysis and contributing to a better understanding of the molecular mechanisms of moulting in copepods. By combining GCN analysis, sample traits and annotation information from public databases we identified relevant modules and hub genes and propose novel candidates with association to moulting and development. For validation, we performed gene knock-down by RNA interference (RNAi) of five genes.

## Methods

### Gene expression data and genome annotation

A normalized gene expression matrix was generated from the RNA-seq data provided by Eichner *et al*. [8], by extracting samples from middle instar ages and old/moulting instar ages of chalimus-1, chalimus-2 and preadult-1 larvae (Figure 1). Transcripts with low expression (not having at least 3 cpm in at least 3 samples) were excluded from the analysis. In this manuscript we are using Ensembl metazoa stable ids (EMLSA[GT]XXXXXXXXXXX) to unanimously identify genes and transcripts in the *L. salmonis salmonis* genome annotation [33]. Gene annotation data were obtained from LiceBase [34].

**Figure 1.**
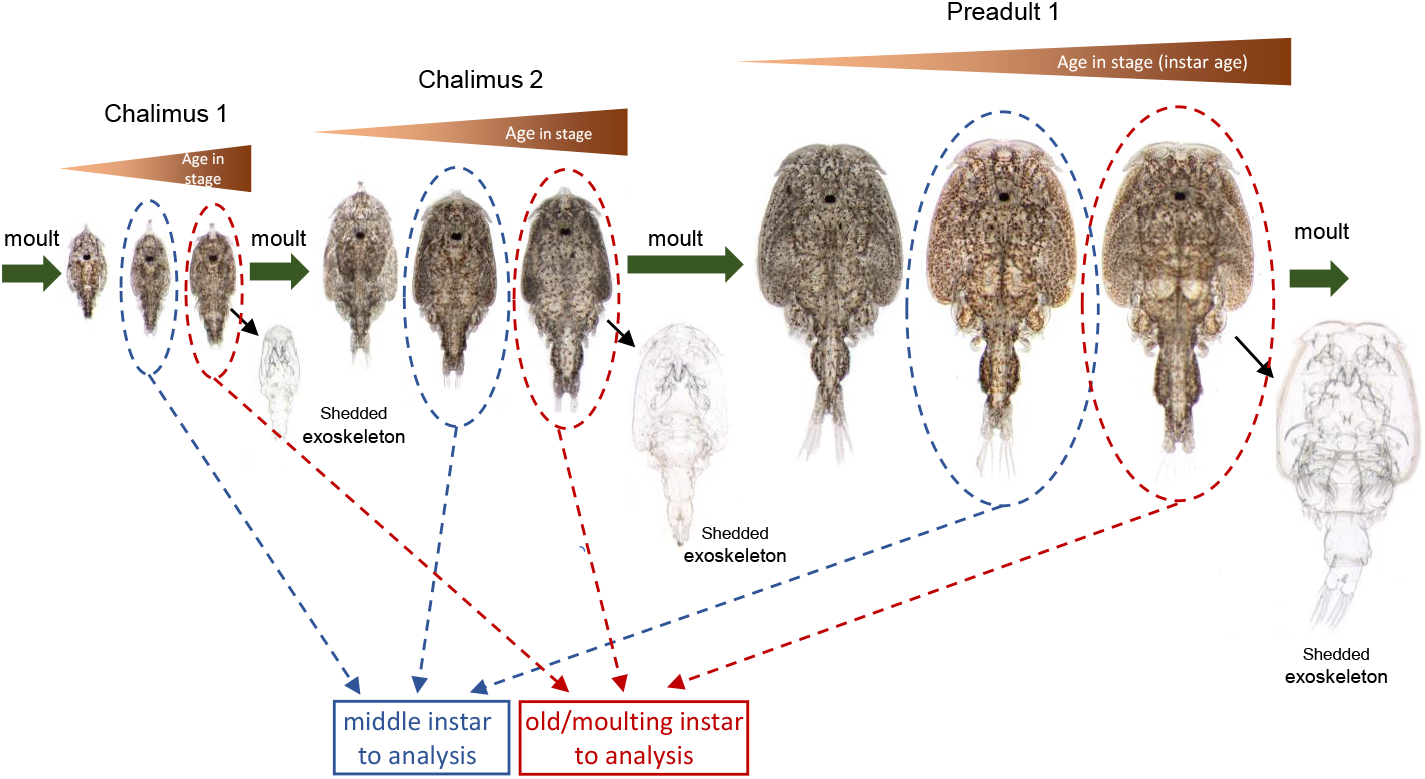
Grouping of sample data and photographs of representative L. salmonis chalimus-1, chalimus-2 and preadult-1 larvae. Within each stage, lice were divided into groups of same instar age: directly after moulting (young), in the middle of the stage (middle) and directly before the moult to the next stage (old/moulting). Moults are represented by a green arrow and a shedded exoskeleton. In this study, data from lice of the middle and old/moulting instar age were used.

### Identification of moulting-associated genes and transcription factor (TF) genes

By combining data from the published literature and LiceBase, we collected genes which are involved in the moulting of salmon lice or known to be associated with the moulting of other arthropods with high confidence. We named these genes as “moulting-associated genes”.

Gene Ontology (GO) annotation information for the salmon louse genes was obtained as previously described [8]. Any salmon louse gene that was annotated by GO terms related to transcription factor (TF) (“GO:0006351”, “GO:0001071”, “GO:0008134”, “GO:0000988” and “GO:0005667”) or child-terms are annotated as TF genes.

### Gene co-expression network (GCN) analysis for identifying important modules and genes associated with moulting and development of salmon louse

In this study, we define the modules and genes that might play a role in the regulation of moulting and development of salmon louse as “important modules” and “important genes”, and we proposed a workflow to identify these important modules and genes based on GCN analysis (Figure 2). Using gene expression profiles, sample traits and gene annotation information as input, this workflow is used to predict the important modules and genes for moulting and development of salmon louse.

**Figure 2.**
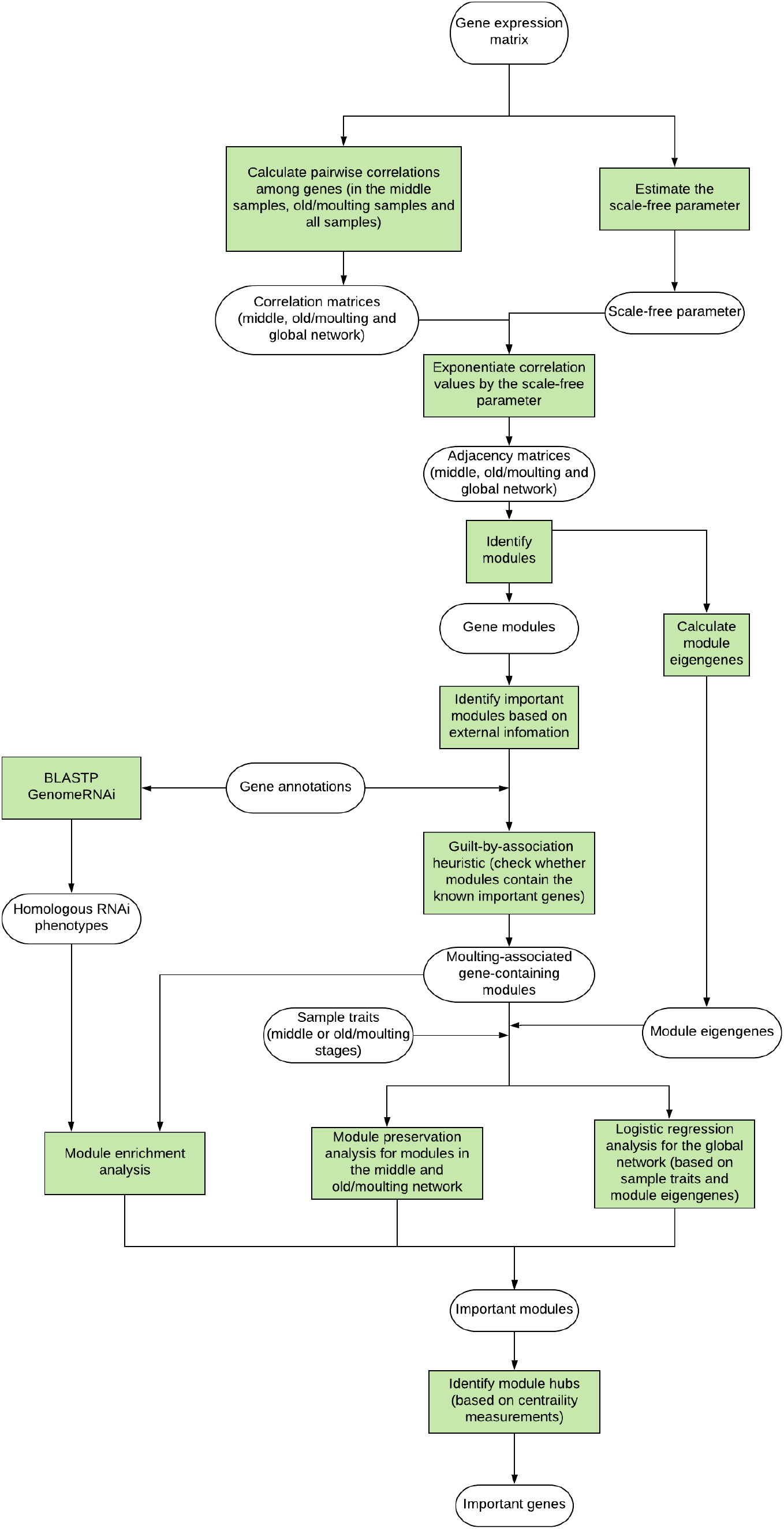
The workflow of gene co-expression network (GCN) analysis for identifying important modules and genes. Methods are highlighted as lightgreen.

### GCN construction, module identification and module eigengene calculation

#### GCN construction and power parameter estimation

GCNs were constructed using the R package WGCNA [35]. A modified version of the biweight midcorrelation (bicor) [36] was adopted to calculate the absolute correlation between pairwise genes (transcripts) (*S*_*ij*_):

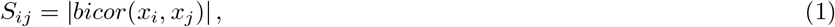

where *x*_*i*_ denotes the expression profile across all samples of transcript *i*. The funnction *bicor* is implemented in the R package WGCNA.

By transforming the correlation by power function, we obtained the adjacency between pairwise transcripts (*A*_*ij*_):

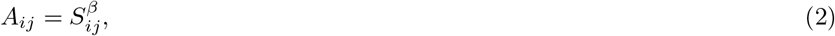

where *β* is the power parameter, and *β* is determined based on whether the corresponding co-expression network exhibits scale-free characteristics and has relatively high connectivities. We chose the suitable power parameter from integers ranging from 1 to 20, by plotting the signed scale-free topology fitting index *R*^2^ against different power parameters, and we also plotted the corresponding network mean connectivity against different power parameters. Details about how the power parameter *β* was estimated can be found in Additional file 1.

With the adjacency matrix *A* we can construct the co-expression network, where each node represents a gene, and the weight possessed by edges between nodes indicates the co-expression relationship between nodes. Although our data is from a transcriptome study we use the term “gene co-expression network” and “eigengene” because transcript quantification was done based on gene-level counts [8].

We constructed three GCNs, based on the gene expression profiles from middle samples, old/moulting samples and all samples (samples from both middle instar ages and old/moulting instar ages).

#### GCN module identification and eigengene calculation

For each GCN, hierarchical clustering was performed for the nodes based on their adjacencies and a dendrogram was obtained. Using this dendrogram as input, a top-down algorithm *cutreeDynamicTree* was applied to identify gene modules. Each module was assigned a unique name as color. For each gene co-expression network, nodes that could not be assigned to any modules were moved to a module called “grey”. The grey module in each of the network was not considered in further analysis.

After identifying modules from each network, a sub-adjacency matrix can be extracted for all the gene members in each module. Then the eigengene for each module was computed as the eigenvector for the largest eigenvalue of the module gene expression matrix by the function *moduleEigengenes* in WGCNA.

### Intramodular centrality measurements and intramodular hub identification

In this study, we adopted three types of centrality measurements to measure the centralities of nodes within each module and identified intramodular hubs.

#### Intramodular Connectivity (kIM)

The connectivity of the *i*th node (*k*_*i*_) in the weighted network is defined as the sum of connection weights between node *i* and the other nodes [37]:

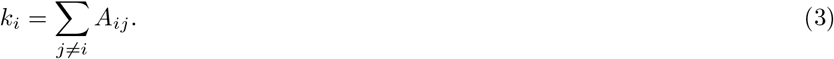

Suppose that there are *Q* modules detected in a network, and they are labeled by *q* = 1, 2, … *Q*, so the connectivity of a node *i* within a module *q* is defined as intramodular connectivity (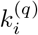 or 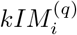):

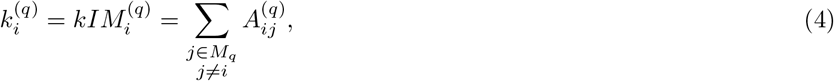

where *M*_*q*_ denotes the set of node indices that correspond to the nodes in module *q*, and *A*^(*q*)^ is the adjacency matrix of module *q*. High intramodular connectivity implies that a node could be a hub within the module.

#### Module membership / module eigengene-based connectivity (kME)

The module membership (or module eigengene-based connectivity) is defined as the value of correlation between module eigengene and the expression profile of the genes (or transcripts) assigned to this module [38]:

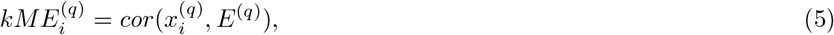

where 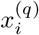 specifies the expression profile in different samples of transcript *i* that is assigned to the module *q*, and *E*^(*q*)^ denotes the eigengene of module *q*.

Since our gene co-expression networks were constructed based on the absolute correlation values between gene expression profiles, we used the absolute value of module membership to measure the centrality of each node within a module:

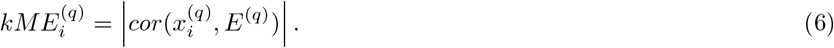

In addition, the module membership of a node for module *q* can be calculated for all nodes in the network:

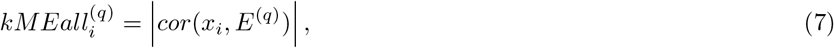

and this definition can be used in the module preservation analysis. The details can be found in Additional file-1.

#### Intramodular weighted betweenness centrality (BC)

The betweenness centrality of a node in an unweighted network (or module) is the number of shortest paths between all other nodes in the network that pass through the node [39]. To calculate the betweenness centralities of nodes in our weighted networks, a generalization of betweenness centrality proposed by Brandes [40] was employed. The approach is implemented in the R package tnet [41].

### Definition of intramodular hubs

We evaluated the centralities of nodes in each module, using intramodular connectivity, absolute module membership and intramodular weighted betweenness centrality. The nodes ranking among the highest ten percent in any of the three centrality measurements of all nodes within a module were defined as intramodular hubs. The node obtaining highest scores in all of the three centrality measurements was defined as “absolute hub”.

Based on the ranks of nodes in three types of centrality measurement, we can calculate the average rank of nodes within each module. Therefore, the absolute hub should have an average rank as 1.

### Module preservation analysis

The preservation of a module between the reference network and a test network can be evaluated based on the alterations in connectivity patterns and density. A well-preservation module in two or more networks should have similar connectivity patterns and nodes in the module should remain being tightly connected. WGCNA provides a series of approaches to evaluate whether a module is preserved and reproducible in another network [42]. In this study, module preservation statistics were computed to compare the two networks constructed based on middle samples and old/moulting samples. For each module preservation statistic, permutation tests were performed to evaluate the significance of the observed value and a *Z* score was obtained. The *Z* scores for all of the module preservation statistics were integrated as a composite summary statistic *Z*_*summary*_. Details about how to calculate module preservation statistics and *Z*_*summary*_ can be found in Additional file 1.

The networks were unsigned, and we set the number of permutation as 200. All the correlations were calculated using the biweight midcorrelation (*bicor*). Modules with a *Z*_*summary*_ smaller than 2 was regarded as non-preserved, while a *Z*_*summary*_ larger than 10 indicated that a module was well preserved across different networks. Since we aimed to identify modules playing a role in the regulation of moulting, the non-preserved modules from the moulting network were of particular interest.

### Regularized logistic regression using module eigengenes as independent variables

We made use of the eigengenes of modules in the global network to perform logistic regression with an elastic-net penalty (*α*=0.5). This task was achieved by setting the binary dependent variable as the label of middle or old/moulting (old/moulting stages were labeled as 1), and using the eigengenes of each module as independent variables.

We used the R package glmnet [43] to perform this analysis, and we adopted the *λ* that gives minimum mean cross-validated error.

### Integrating information from external databases and enrichment analysis

Data from FlyBase [44, 45] and GenomeRNAi [46] were extracted and used to identify homologous observable phenotypes and lethal phenotypes enriched modules. To detect homologous sequences in *Drosophila melanogaster*, we ran BLASTP with E-value cutoff as 1*e* − 10 on the corresponding protein sequences of salmon louse transcripts against protein sequences from *Drosophila*. Only best hits were considered. After mapping the protein IDs of the homologues from *Drosophila* to gene IDs, RNAi knock-down phenotype information were mapped to data from GenomeRNAi. If a salmon louse protein had more than one *Drosophila* homologue with identical maximum bitscore, all the homologues were used to search for RNAi phenotypes. BLASTP searches of all salmon louse predicted amino-acid sequences were performed to find paralogues.

### Enrichment analysis of modules

Based on the GO annotation file for salmon louse genes from LiceBase, GO enrichment analyses were performed for each modules identified in the middle, moulting and global network using the fisher statistic and “elim” algorithm provided by R package topGO [47].

Furthermore, with the information from the *Drosophila* homologues-based transcript-phenotype list (Additional file 2-Table S1), we conducted two enrichment analyses for each module identified in all networks. The p-values of these enrichment analyses were obtained based on hypergeometric tests, to determine whether transcripts with homologue observable phenotypes or homologue lethal phenotypes in *Drosophila* were significantly enriched within a module. Based on the suggestions from [48], we used the raw p-values of our enrichment analyses, and the cutoff of p-values was set as 0.05 for all the enrichment analyses.

### Selecting important modules for further analyses

We were interested in identifying gene modules which are likely to play a role in the moulting and development of salmon louse, and we chose important modules based on three analyses: the module preservation analysis, the regularized logistic regression analysis and the *Drosophila* homologues-based enrichment analysis. According to the guilt-by-association (GBA) heuristic [49], nodes in the moulting-associated transcripts-containing modules are more likely to play a role in the moulting and development of salmon louse, and we conducted a focused search among modules containing at least one known moulting-associated transcript. Therefore, moulting-associated transcripts-containing modules satisfying any of the following criteria were chosen for further studies: 1) non-preserved modules in the moulting network (*Z*_*summary*_ < 2); 2) the eigengenes of modules from the global network obtained positive coefficients from the regularized logistic regression analysis (the module with largest coefficient value should be prioritized); 3) modules that are significantly enriched by transcripts with observable and lethal RNAi phenotypes from homologues (*p* − *value* < 0.05) (Figure 2).

### Selecting important genes as knock-down candidates from important modules

Since many researchers have proposed that hubs in a biological network tend to be more important [50–52], we chose RNAi knock-down candidates among the hubs of the important modules. For each selected module, we gave prime consideration to the absolute hub. If no absolute hub was detected, knock-down candidates were chosen from other intramodular hubs. Hubs with less paralogues and little annotation information were then given priority.

### Differential gene expression (DE) analysis and regulatory impact factor (RIF) calculation

We calculated the regulatory impact factors (RIF) for all the transcripts annotated as TF, based on the metric proposed in [53].

The first step was to perform differentially expressed (DE) analysis to compare the middle group and the old/moulting group, and the statistics used to test the null hypotheses were calculated based on standardized rank-sum Wilcoxon test. We computed the permutation adjusted p-values using the step-down maxT multiple testing procedures, which provide strong control of the family-wise Type I error rate (FWER). The functions are implemented in the R package multtest [54]. Transcripts with an adjusted p-value smaller than 0.05 were identified as differentially expressed between middle instar ages and old/moulting instar ages.

The first RIF value (*RIF* 1) for *f* th TF transcript was defined as:

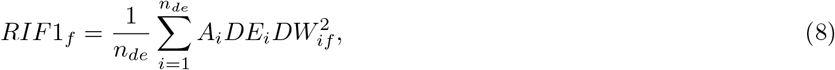

where *n*_*de*_ is the number of DE transcripts; *A*_*i*_ represents the average expression of *i*th DE transcript across the two groups, and *DE*_*i*_ is the statistics obtained from the previous DE analysis. *DW*_*if*_ is the abbreviation for differential wiring, which means the change of correlation between *f* th TF and the *i*th DE transcript across the two groups:

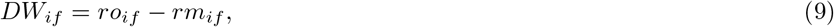

where *ro*_*if*_ and *rm*_*if*_ are the co-expression correlation between *f* th TF and the *i*th DE transcript in the old/moulting samples and the middle samples, respectively. The second RIF value (*RIF* 2) was computed as followed:

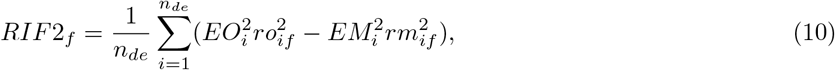

where 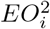 and 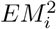 denote the square of the average expression value of the *i*th DE transcript in the old/moulting samples and the middle samples;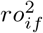 and 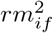 are the square of the co-expression correlation between *f* th TF and the *i*th DE transcript in the old/moulting samples and the middle samples.

### RNA interference experiments

#### RNA interference

Double-stranded RNA (dsRNA) was produced using MEGAscript ® RNAi Kit (Ambion) according to supplier’s instructions using following primers added the T7 sequence (TAATACGACTCACTATAGGGAGA): EMLSAG00000001458: for-ward (fw)_1458: CAAGCTGTTATTGATTGGCGATTC, revers (rv)_1458: CG GCATATTTAACTGATCAGCGTA; EMLSAG00000003179:fw_3179: GCGTAAAAGTTGCGTACAATCTGA, rv_3179: GTTTATTGGGTGTGATGAATCCGA; EML-SAG00000005299: fw_5299: GTATGATGACGGACATGCTCAAGG, rv_5299: GGCCTGTTTATAGTCGGTAGCCAT; EMLSAG00000004347: fw_4347: AAACGGCG CGAGGAGGTGAATA, rv_4347: GGTGGGTTTCTTTCCTGGCTTGTT; EML-SAG00000008959: fw_8959: GCCTCCGGTTCGGATGAAGAA, rv 8959: AGGATC AGAGGGGCCACAAGTGTC.

#### RNA interference on nauplia and fish challenge with the emerging copepodids

RNA interference was conducted in nauplia as described in [55], but with 2 *µg* fragment. Infection was done in single tanks with 60 copepodids per fish on three fish for each fragment and for the control as described in [56] (10°C, full salinity). The remaining copepodids were stored on RNAlater® (Invitrogen) for later measurement of transcript down regulation. Sampling was done after 16 days when lice from control group were in chalimus or preadult-1 stage. All lice were sampled from fish and photographs were taken under the binocular in a drop of seawater with a cover slide on top. Number of lice and size measurements on photographs were recorded. Genes to be knocked down were: EMLSAG00000001458, EMLSAG00000003179 and EML-SAG00000005299. A second trial with knock-down larvae of EMLSAG00000001458 with 100 copepodids on three fish each was done. The outflow water from the tanks was filtered and lice in the flow out were counted two hours after infection and 24 hours after infection. One fish from knock-down and control group each were terminated three days after infection and lice were fixed on Karnovsky’s fixative for histological investigation. The other fish were terminated eight days after infection.

#### RNA interference on preadult lice

RNA interference in preadult-2 lice was done as described in [57]. In short, the fragment was injected into preadult-2 lice, which were put on fish again until most of the control lice showed its second pair of egg strings. All lice were sampled and photographed. The lice egg strings were laid into single flow through wells for hatching observation. Lice were either stored on RNAlater® for qPCR measurements or on Karnovski’s reagent for histological investigation. Three different experiments were conducted, one with double-stranded RNA for EMLSAG00000001458, EML-SAG00000005299 and EMLSAG00000003179 as well as a control (35 days), one with dsRNA for EMLSAG00000004347 and a control (40 days) and one with dsRNA for EMLSAG00000008959 and a control (37 days). The cod trypsin RNA was used as a non-target control fragment [57].

#### RNA extraction, cDNA synthesis and qPCR measurements

Nauplia were divided into five (or four in case of control group from second infection trial) batches of 30 to 40 nauplia. RNA from nauplia was extracted by a combination of Trizol and RNAeasy micro kit as previously described [55]. RNA was frozen at -80 °C until usage. cDNA synthesis was conducted using the AffinityScript QPCR cDNA Synthesis Kit (Agilent) according to suppliers recommendations. Gene expression of the target gene was measured by quantitative real-time PCR in control and knock-down group. QPCR was carried out in duplicates using the salmon louse elongation factor 1*α* [58] as well as Adenine Nucleotide Translocator 3 (ADT3) [56] (in nauplia only) as a standard. SYBR® Green PCR master mix and the following primers were used: forward (fw)_1458: CAAGCTTCTTGTGGGAAACAAATG, reverse (rv)_1458: GCTGCCATAGTCATAAAAGCTTGC; fw_3179: GGTTCGGATTCATCACACCCA, rv_3179: CGTGGAGGGATAGGACAAACTTTG; fw_5299: GTTCTCG GAGAAAATAATGTTGCG, rv_5299: TGTTCAGTGATTTCCAGTGCTTCC; fw_8959: CCATCAATTTCCAAGTGGAGGA, rv_8959: CCTTCGCATATCTCTTCCTCTTCA; fw 4347: GTCTGGACAAGGAGAAAATTCTGC, rv_4347: CTC CGTGGTTTTGGCATTGA; fw_LsEF1*α*: GGTCGACAGACGTACTGGTAAATC C, rv_LsEF1*α*: TGCGGCCTTGGTGGTGGTTC, fw_LsADT3: CTGGAGAGGGA ATTTGGCTAACGTG, rv_LsADT3: GACCCTGGACACCGTCAGACTTCA. Applied Biosystems 7500 Fast Real-Time PCR system was used for thermal cycling and quantification in 10 *µl* reactions (initiation 50 °C 2 min, 95 °C 2 min, then 40 cycles of 95 °C 15 seconds, 60 °C 1 min). A melting curve 60 °C to 90 °C was performed. Relative gene expression was calculated using the differences in threshold cycle (CT) between gene of interest and standard genes.

#### Histology

Lice were prepared as described in [59] for histological investigation. For plastic embedding, lice were washed twice in phosphate buffered saline, dehydrated in a graded ethanol series, pre-infiltrated with Technovit/ethanol (50/50) for four hours (Technovit 7100, Heraeus Kulzer Technique) and infiltrated with Technovit and hardener overnight. Two micrometre thick sections were cut with a microtome (Leica RM 2165) and stained with toluidine blue (1% in 2% borax) for one minute. The stained sections were mounted using Mountex (Histolab Products).

## Results

### Identification of moulting-associated and TF genes

Among the transcripts in our RNA-seq data, we found 40 moulting-associated transcripts and 32 of them were retained after low expression filtering. The list of moulting-associated transcripts and the relevant publications can be found in Table 1.

**Table 1.** ID, annotation and relevant publications for the moulting-associated transcripts

| TranscriptID | Annotation from LiceBase | Publication |
| --- | --- | --- |
| EMLSAT00000012651 | Nuclear receptor subfamily 1 group I member 2 | [92] |
| EMLSAT00000004055 | Phosphoacetylglucosamine mutase | [93] |
| EMLSAT00000002853 | Chitin synthase 8 | [17] |
| EMLSAT00000007308 | Chitin synthase 8 | [94] |
| EMLSAT00000012864 | Probable glucosamine 6-phosphate N-acetyltransferase | [95] |
| EMLSAT00000000683 | Glutamine-fructose-6-phosphate aminotransferase [isomerizing] 2 | [96] |
| EMLSAT00000008812 | Acidic mammalian chitinase | [17] |
| EMLSAT00000005289 | Probable chitinase 2 | [17] |
| EMLSAT00000008235 | Acidic mammalian chitinase | [17] |
| EMLSAT00000005464 | Chitooligosaccharidolytic beta-N-acetylglucosaminidase | [97] |
| EMLSAT00000008487 | Beta-hexosaminidase subunit alpha | [98] |
| EMLSAT00000000150 | N-acetylglucosamine-6-phosphate deacetylase | [16] |
| EMLSAT00000001322 | Glucosamine-6-phosphate isomerase | [16] |
| EMLSAT000000004643 | Hexokinase type 2 | [99] |
| EMLSAT00000005083 | Hexokinase type 2 | [99] |
| EMLSAT00000001264 | Hexokinase type 2 | [99] |
| EMLSAT00000008931 | Glucose-6-phosphate isomerase | [100] |
| EMLSAT00000003220 | Mannose-6-phosphate isomerase | [101] |
| EMLSAT00000007185 | Phosphoglucosmutase-2 | [101] |
| EMLSAT00000011599 | Phosphoglucosmutase-1 | [101] |
| EMLSAT000000001990 | TMEM47 | [102] |
| EMLSAT00000001904 | Ecdysone-induced protein 78C | [103] |
| EMLSAT00000000733 | Probable nuclear hormone receptor HR3 | [104] |
| EMLSAT000000008543 | Hormone receptor 4 | [104] |
| EMLSAT000000008902 | Nuclear hormone receptor FTZ-F1 | [105] |
| EMLSAT00000008651 | Ecdysone-induced protein 74EF isoform A | [92] |
| EMLSAT00000000188 | Zinc finger protein 142 | [106] |
| EMLSAT000000009170 | Growth hormone secretagogue receptor type 1 | [55] |
| EMLSAT00000008881 | Cathepsin L | [107] |
| EMLSAT00000001278 | Ion transport peptide | [108] |
| EMLSAT00000008115 | CHH-like protein | [109] |
| EMLSAT00000001150 | Cytochrome P450 307a1 | [69] |

There were 433 transcripts annotated as TF, and 231 TF were retained after low expression filtering (Additional file 2-Table S8).

### GCN construction and module identification

To detect the genes that might be involved in the regulation of moulting and development, we analyzed the RNA-seq data sampled from the middle instar ages and old/moulting instar ages of chalimus-1, chalimus-2 and preadult-1 larvae (Figure 1). Of the 45 samples, 18 samples were from middle instar ages, and 27 samples were from old/moulting instar ages. After filtering transcripts with low expressions, 7108 transcripts were retained for network analysis.

Three GCNs were constructed for different sample groups. The first GCN was generated using all samples labeled as “middle instar age”. Meanwhile, an “old/moulting instar age” GCN was created for all old/moulting samples. To further facilitate our analysis, a GCN based on all samples from middle and old/moulting instar ages was built. The three GCNs were thus denoted as middle network, moulting network, and global network, respectively.

We set the power parameter *β* as 7 to make sure that the networks satisfied scale-free topology approximately while having relatively high mean connectivities (Figure 3). The adjacency matrices of the three networks can be found in Additional file 7.

**Figure 3.**
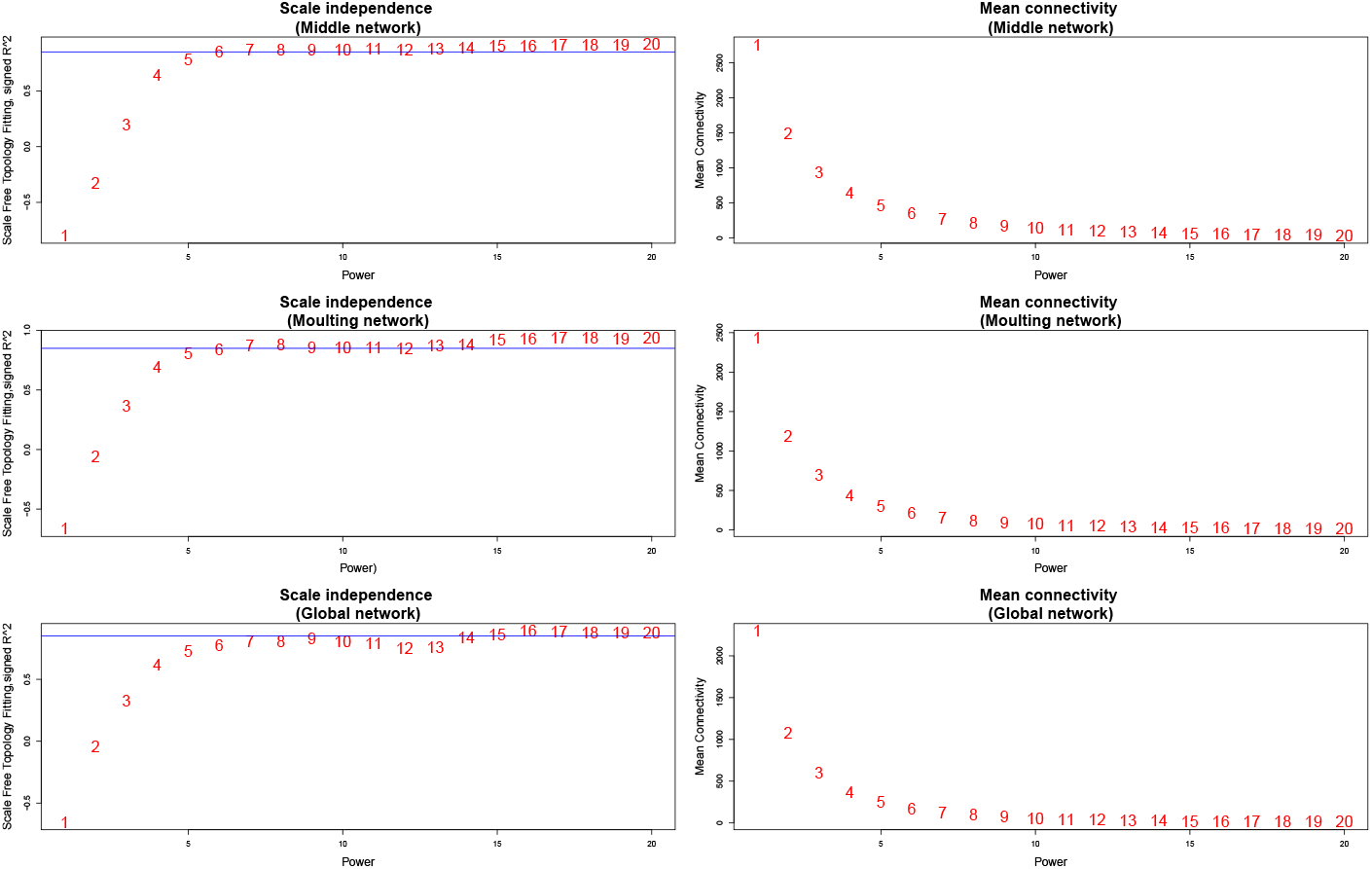
Network scale-free topology and mean connectivity for different power parameters ranging from 1 to 20. Plots on the left show the scale-free topology fitting index (y-axis) as a function of the power (x-axis) in the middle, moulting, and global network, respectively. A horizontal line is added at y=0.85 in each plot. Plots on the right show the mean connectivity (y-axis) as a function of the power (x-axis) in the middle, moulting, and global network, respectively.

In the module identification process, 83, 60 and 78 modules were found in the middle, moulting and global network, respectively, and the module sizes ranged from 32 to 333. There were 203, 444 and 506 genes assigned to the grey module of the middle, moulting and global network, respectively (Additional file 2-Table S2-S4). Genes in the grey modules were not considered for further analysis. Dendrogram and module identification results for the three networks were displayed in Additional file 3-Figure S1. Eigengenes for each module in all the three networks were also obtained.

### The centralities and distribution of moulting-associated genes across modules

To preliminarily examine the essentiality of intramodular hubs, we calculated the centralities for the 32 moulting-associated transcripts. Based on our definition of intramodular hubs, there were 6, 12 and 8 moulting-associated transcripts identified as intramodular hubs in the middle, moulting and global network, respectively (Additional file 2-Table S5-S7). The transcript EMLSAT00000005083 (annotated to encode hexokinase type 2, Table 1) was identified as intramodular hub in all the three networks, and it was the absolute hub in the module “lightcyan1” of the global network.

We examined how the 32 moulting-associated transcripts were distributed across modules in the three networks. There were 25, 20 and 24 modules containing moulting-associated transcripts in the middle, moulting and global network, accounting for 30.1%, 33.3% and 30.8% in the three networks, respectively. The numbers of moulting-associated transcripts in these modules ranged from 1 to 4 (Additional file 2-Table S2-S4).

### Module preservation analysis

To identify genes which may play a role in the moulting of salmon lice, we detected non-preserved modules from the moulting network based on module preservation analysis. Five modules from the moulting network were found as non-preserved, and the module sizes ranged from 41 to 100 (Additional file 2-table S3). Strong correlations among genes in these non-preserved modules were only observed in the moulting network, and two non-preserved modules (yellowgreen and lavenderblush3) contained moulting-associated transcripts (EMLSAT00000008812 and EMLSAT00000012651) (Additional file 2-Table S6). Notably, the moulting-associated transcripts were also identified as intramodular hubs in these modules. The transcript EMLSAT00000008812 (annotated to encode chitinase, Table 1) was ranked eighth (based on connectivity) in the “yellowgreen” module, and the transcript EMLSAT00000012651 (annotated as EcR, Table 1) was ranked third (based on betweenness centrality) in the “lavenderblush3” module. We thus hypothesized that transcripts in these two modules could be important for salmon louse moulting, and hubs from these modules should be considered as important.

Eight modules from the middle network were identified as non-preserved, and the module sizes ranged from 54 to 109 (Additional file 2-Table S2). Three non-preserved modules (darkseagreen4, brown4 and lightcyan1) were found containing one moulting-associated transcript (Additional file 2-Table S5). However, none of these moulting-associated transcripts were intramodular hubs in the middle network.

### Regularized logistic regression analysis on the global co-expression network

To compare the intramodular overall gene pexression patterns between the middle sample group and old/moulting sample group, we performed elastic net regularization-based logistic regression using the eigengenes of module from the global network as independent variables. As a result, we found modules with eigengenes that were highly expressed in one sample group but lowly expressed in the other sample group. From the 78 module eigengenes, we identified 15 eigengenes with non-zero coefficient (ranging from -1.75 to 0.963), and six of the 15 corresponding modules contained one known moulting-associated transcripts (Additional file 2-Table S4). It was noteworthy that module “steelblue” possessed the largest positive coefficient and contained one moulting-associated transcript (EMLSAT00000001150) as intramodular hub, which was ranked second in the betweenness centrality measurement (Additional file 2-Table S7).

When checking the absolute value of regression coefficients, three modules (magenta, lightcyan and ivory) were found with absolute coefficient larger than 1. The moulting-associated transcripts were found in two of the three modules (lightcyan and ivory). The coefficients of all the three modules were negative, indicating that genes in these modules exhibited much higher expressions in middle samples. Notably, two modules (indianred4 and lavenderblush3) with negative regression coefficients contained moulting-associated transcripts (EMLSAT00000000733 and EMLSAT00000008543) annotated as hormone receptor 3 (Hr3) and hormone receptor 4 (Hr4) (Table 1). In the module “indianred4”, EMLSAT00000000733 was ranked eighth in the connectivity measurement.

Differentially expressed transcripts between middle group and old/moulting group were found in all the modules with non-zero regression coefficients, and the proportions ranged from 36.9% to 98.4% (Additional file 2-Table S4).

### Integrating information from external databases

We identified homologous genes in *Drosophila melanogaster* for the salmon louse transcripts and then searched for RNAi phenotypes for these homologues in the GenomeRNAi database. We found homologous RNAi phenotypes for 3059 salmon louse transcripts. Additional file 2-Table S1 maps salmon louse transcripts to the RNAi phenotypes of the corresponding homologues in *Drosophila*.

### Enrichment analysis of modules

Based on the GO annotation file for the salmon louse transcripts, we performed GO enrichment analysis for all the modules to preliminarily elucidate the biological functions of the modules. The GO term with smallest p-value in each category (Biological Process(BP), Molecular Function(MF) and Cellular Component(CC)) were recorded (Additional file 2-Table S2-S4).

To further identify modules which are more likely to contain important genes for lice development, we conducted enrichment analyses for all the modules based on the homologues-based transcript-phenotype list. The transcripts with observable RNAi phenotypes were significantly enriched in 16, 13, and 14 modules in the middle, moulting and global network, accounting for 19.3%, 21.7%, and 17.9% in total modules, respectively. Analogously, 14, 14, and 9 modules were detected as enriched by transcripts with lethal RNAi phenotypes in the middle, moulting and global network, accounting for 16.9%, 23.3% and 11.5% in total modules. We found a relatively large overlap between the two enrichment analyses: 10, 11 and 7 modules (accounting for 12.0%, 18.3% and 9.0% in total modules) were identified as being significantly enriched by both observable and lethal RNAi phenotypes in the middle, moulting and global network (Additional file 2-Table S2-S4).

### Differential gene expression (DE) analysis and regulatory impact factor (RIF) calculation

All 45 samples were divided into middle and old/moulting groups to find DE transcripts. There were 1364 transcripts differentially expressed between the two groups. The list of DE transcripts facilitated calculation of the RIF scores for all transcripts with GO annotation as TF.

For the 231 TF transcripts, RIF scores were computed (Additional file 2-Table S8). It is noteworthy that EMLSAT00000003849 (annotated as forkhead box protein O(FOXO)) obtained highest RIF scores from both methods. This transcript is also an intramodular hub of a moulting-associated transcripts-containing module in both of the middle and moulting network.

### Selecting important modules for further analyses

In the module preservation analysis, two modules (yellowgreen and lavenderblush3) from the moulting network were detected based on our criteria. In the regularized logistic regression analysis, two modules (steelblue and green) from the global network passed the criteria. In the homologues-based enrichment analysis, one (mediumpurple3), two (darkolivegreen and violet) and one (turquoise) module were found satisfying the criteria from the middle, moulting and global network, respectively. In summary, one, four and three modules from the middle, moulting and global network were selected for further analyses.

### Examining intramodular hubs and selecting important genes as knock-down candidates from important modules

After determining important modules, we investigated the hubs of these modules to understand their roles in moulting and development of salmon louse. For each of the eight chosen modules, we examined their hub with highest average rank (Additional file 2-Table S12). The absolute hubs (EMLSAT00000003844 and EMLSAT00000001458) of the two non-preserved modules (lavenderblush3 and yellow-green) selected from the moulting network are annotated as epithelial cell transforming 2 (ECT2) and Ras-related protein Rab-1A (RAB1A), respectively. For the four modules selected from the enrichment analysis, EMLSAT00000000929, EMLSAT00000005299, and EMLSAT00000012769 were identified as absolute hubs, annotated as high density lipoprotein-binding protein (HDLBP), ER membrane protein complex subunit 3 (EMC3) and laminin subunit beta-1 (LanB1). EMLSAT00000010555 had the highest average rank in the module (turquoise) from the global network, annotated as stress-induced-phosphoprotein 1 (STIP1). No absolute hubs were found in the two global modules selected from the regression analysis, and the nodes with highest average rank in these modules were EMLSAT00000007421 and EMLSAT00000012693, both of them were identified as differentially expressed transcripts between the middle group and old/moulting group. EMLSAT00000007421 was annotated as cuticular protein 62Bb (Cpr62Bb), and few annotation was found for the hub EMLSAT00000012693.

To validate the importance of genes in the selected modules in moulting and development of salmon louse within the limited accesses to RNAi experiments, we selected RNAi knock-down candidates from three important modules. Since the important modules were selected based on three analyses, we selected one module from each of the three analyses. Firstly, we choose the module “yellowgreen” from the two non-preserved modules (the other one is lavenderblush3) in the moulting network. The module “yellowgreen” had larger size then the module “lavenderblush3”, and the absolute hub of the module “yellowgreen” had higher score of the absolute module membership. From the two moulting modules (darkolivegreen and violet) selected from the enrichment analysis, we chose the module “violet” for further analysis, since it contained more transcripts annotated as TF, and these transcripts obtained higher scores in the regulatory impact factor analysis than those found in the module “darkolivegreen”. Furthermore, with regards to the proportion of transcripts annotated as TF, the module “violet” and “yellowgreen” ranked as first and third among all modules in the moulting network. Finally, we selected the module “steelblue” in the global network for further analysis, because the eigengene of this module obtained largest coefficient in the regularized logistic regression analysis. Details of the three selected modules can be found in Table 2 and Additional file 2. For the module “yellowgreen” and “violet” from the moulting network, we chose the absolute hub for RNAi experiment. According to the criteria discussed in the method section, we chose another one hub without paralogues from each module to knock down.

**Table 2.** Information on selected modules for selection of knock-down candidates.

| Module | Network | Size | $Z_{summary}$ | Number of moulting-associated genes | Number of TF genes | $p - value$ | |
| --- | --- | --- | --- | --- | --- | --- | --- |
|  |  |  |  |  |  | RNAi phenotype | RNAi lethal phenotype |
| yellowgreen | moulting | 81 | 1.92 | 1 | 7 | 0.84 | 0.54 |
| violet | moulting | 83 | 8.52 | 1 | 10 | 0.026 | 0.0076 |
| steelblue | global | 79 | — | 1 | 0 | 0.90 | 0.94 |

No absolute hub was found in the module “steelblue” from the global network, and the hub (EMLSAT00000007421) with highest average rank was annotated to encode cuticle protein. Among the 12 intramodular hubs found in the module “steelblue”, three (EMLSAT00000012111, EMLSAT00000008158 and EMLSAT00000012113) were annotated to encode proteins with the chitin binding peritrophin-A domain (PF01607); four (EMLSAT00000007421, EMLSAT00000007422, EMLSAT00000009987, and EMLSAT00000010209) were annotated to encode cuticle proteins (PF00379); one (EMLSAT00000004870) was annotated to encode protein with the polyprenyl synthetase domain (PF00348), and the moulting-associated transcript (EMLSAT00000001150) was annotated to encode cytochrome P450 (PF00067) (Additional file 2-Table S10, Table 1). Among the three hubs with few annotation information, we chose one (EMLSAT00000004347) with least number of paralogues to knock down. The details of all the knock-down candidates were collected in Table 3.

**Table 3.**
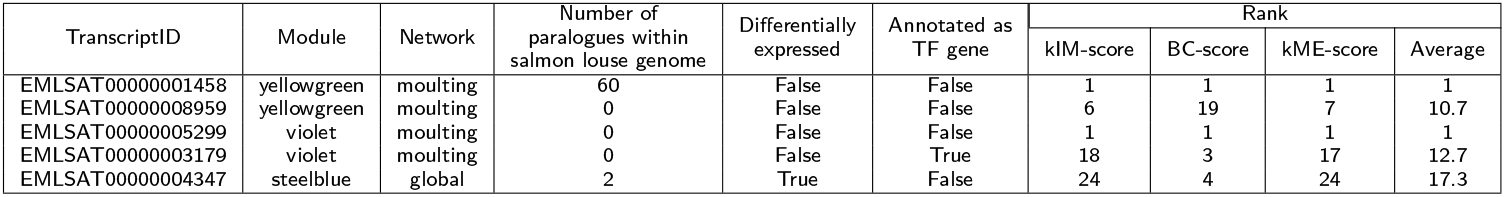
Information of the selected knock-down candidate genes.

| TranscriptID | Module | Network | Number of paralogs within salmon louse genome | Differentially expressed | Annotated as TF gene | Rank |  |  |  |
| --- | --- | --- | --- | --- | --- | --- | --- | --- | --- |
|  |  |  |  |  |  | kIM-score | BC-score | kME-score | Average |
| EMLSAT00000001458 | yellowgreen | moulting | 60 | False | False | 1 | 1 | 1 | 1 |
| EMLSAT00000008959 | yellowgreen | moulting | 0 | False | False | 6 | 19 | 7 | 10.7 |
| EMLSAT00000005299 | violet | moulting | 0 | False | False | 1 | 1 | 1 | 1 |
| EMLSAT00000003179 | violet | moulting | 0 | False | True | 18 | 3 | 17 | 12.7 |
| EMLSAT00000004347 | steelblue | global | 2 | True | False | 24 | 4 | 24 | 17.3 |

### RNA interference on nauplia and infection of salmon with the emerging copepodids

Measurement of gene expression in copepodids before infection showed down regulation of all targeted genes (t-test: *p* − *value* < 0.05) with varying knock-down efficiency. For genes EMLSAG00000001458, EMLSAG00000005299 and EMLSAG00000003179, efficiency was 94%, 84% and 89%, respectively. At termination after 16 days no lice were found on the fish infected with copepodids from EMLSAG00000001458-KD group (Figure 4, left panel). There was no significant difference in the number of lice between control group and EMLSAG00000005299-knock-down (KD) group as well as EMLSAG00000003179-KD group, and the development of lice from all groups found on fish was similar (Figure 4, right panel). No difference in the phenotype could be observed under the binocular or by size measurements.

**Figure 4.**
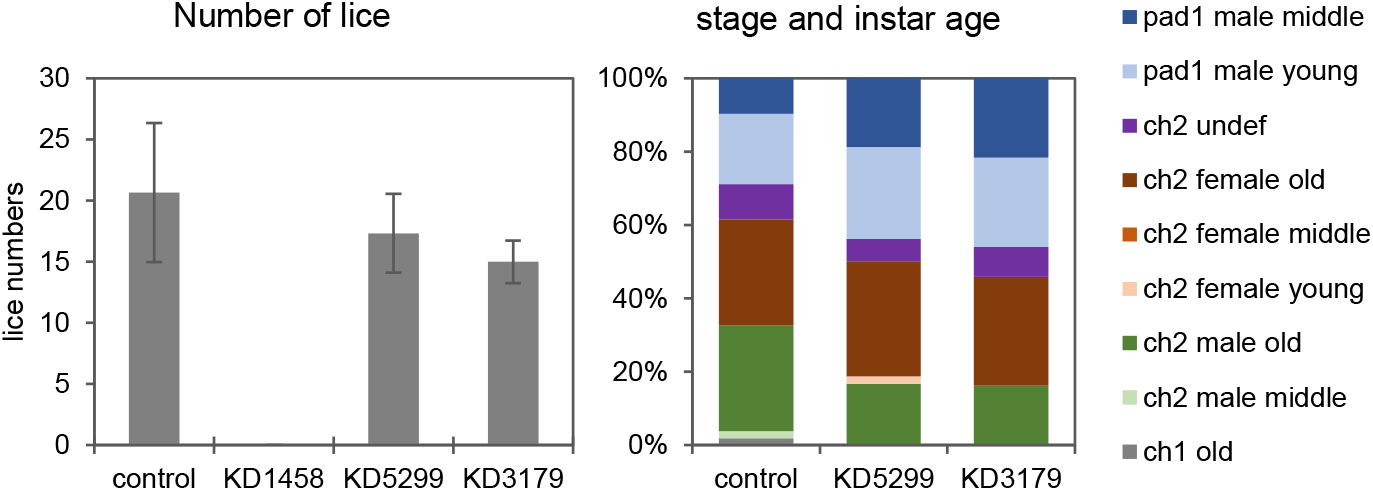
Number of L. salmonis per fish (left panel) and distribution of stage and instar age (right panel). Lice were collected at termination of the knock down experiment, 16 days post infection, dsRNA was introduced in nauplia larvae. pad= preadult, ch= chalimus. Sample names represent dsRNA target transcripts; KD1458 : EMLSAT00000001458, KD5299 : EMLSAT00000005299, KD3179 : EMLSAT00000003179. Error bars represent + − 1 standard deviation, N=3

#### Second trial with EMLSAG00000001458 knock-down

Since no lice with EMLSAG00000001458 knocked down were found at termination after 16 days on the fish, we were interested in finding out whether this could be due to reduced infection success or due to problems with development and moulting. A second infection trial for qualitative measurement was done. Knock down efficiency measured in copepodids before infection was 95%. After two hours, 30, 37 and 33 lice were found in the filtered flow through water of tanks from fish of the control group, and 32, 57 and 35 lice were found from tanks of the knock-down group. After 24 hours, 9, 9 and 4 lice were found in the flow out from control fish, and 9, 8 and 4 lice were found from knock-down fish. No lice were found in the filters after three days. At termination of the first fish at day three after infection there were 10 lice on the control fish and 14 on the knock-down fish. These were sampled for histological investigation. No differences were observed in the histological sections (Additonal file 3-Figure S3). Eight days after infection, lice had developed to chalimus-1 on control fish (13 on one fish, 39 on the other), but no lice were found on one of the fish with knock-down samples and two copepodids on the other fish.

#### Knock-down in preadult lice

At sampling lice were in the adult stage. Down regulation was on average 77% for EMLSAG00000001458-KD group, 47% for EMLSAG00000003179-KD and 68% for EMLSAG00000004347-KD group. Lice from EMLSAG00000001458-KD group and EMLSAG00000008959-KD group had no egg string. Length measurements for body parameters (cephalothorax and genital segment length) as well as egg strings are shown in Table 4. Egg strings of all groups with egg strings present hatched and produced viable normal looking offspring. Histological sections were done for EMLSAG00000001458-KD and EMLSAG00000004347-KD lice. Histology of different tissues is shown in Additonal file 3-Figure S2. EMLSAG00000001458 knock-down lice did not develop normal looking oocytes (Additonal file 3-Figure S2 f) and the ovaries did not contain any oogonia (Additonal file 3-Figure S2 j). The cellular structures of the subcuticular tissue of the cephalothorax were changed and only

**Table 4.**
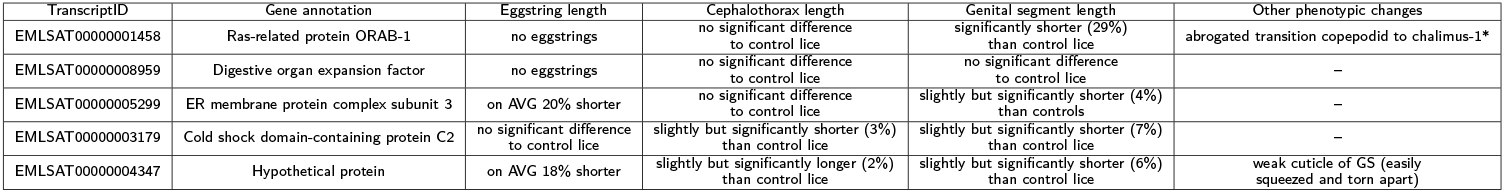
Observed phenotypes by RNAi for the selected knock-down candidates. All phenotypes were assessed in adult female lice after injection of dsRNA at the pre-adult 2 stage except for *: phenotypic change was observed at larval stages after dsRNA treatment of nauplii (see Figure 4).

loose connection between cells was observed (Additonal file 3-Figure S2 n), while the subcuticular tissue of the genital segment seemed not to be affected in the same way. At sampling EMLSAG00000004347 knock-down lice showed a weak genital segment, which was easily squeezed and teared apart when handling the lice. In the histological sections, the cuticle and subcuticular tissue of the genital segment (Additonal file 3-Figure S2 s) did not show obvious differences to the control louse.

### Examining the RNA interference experiments data from LiceBase

From LiceBase, RNAi experiments for 188 genes were collected, and 112 genes among them appeared in our three networks. 10 genes in six of the eight selected modules were found with observable RNAi phenotypes (including the RNAi experiments results from this study). One gene from the selected module “darkolivegreen” had been knocked down, but no phenotype was observed. No RNAi results were found for the genes in the selected module “lavenderblush3” (Additional file 2-Table S13).

Notably, one hub (EMLSAG00000009839) from one non-preserved module (sky-blue) of the moulting network show reduced survival in the RNAi experiments, although this module did not contain any known moulting-associated genes. The absolute hub (EMLSAG00000005382) of the module “blue2” in the middle network show shorter eggstrings in RNAi experiments, and this module contained two known moulting-associated genes.

We also found RNAi experiment records of four genes (EMLSAG00000010968, EMLSAG00000006642, EMLSAG00000007048 and EMLSAG00000004159) which obtained highest average rank in four modules. These four modules did not satisfy any of the criteria for being an important module for salmon louse moulting and development, and the four genes did not show any observable phenotype in RNAi experiments (Additional file 2-Table S13).

### Examining the modules where the RNAi candidates were selected

In the RNAi experiments, all five selected gene candidates show observable phenotypes, and we thus examined the three modules they were from. We plotted heatmaps of scaled gene expression profiles and barplot of scaled eigengene expression for each of the three module (Figure 5).

**Figure 5.**
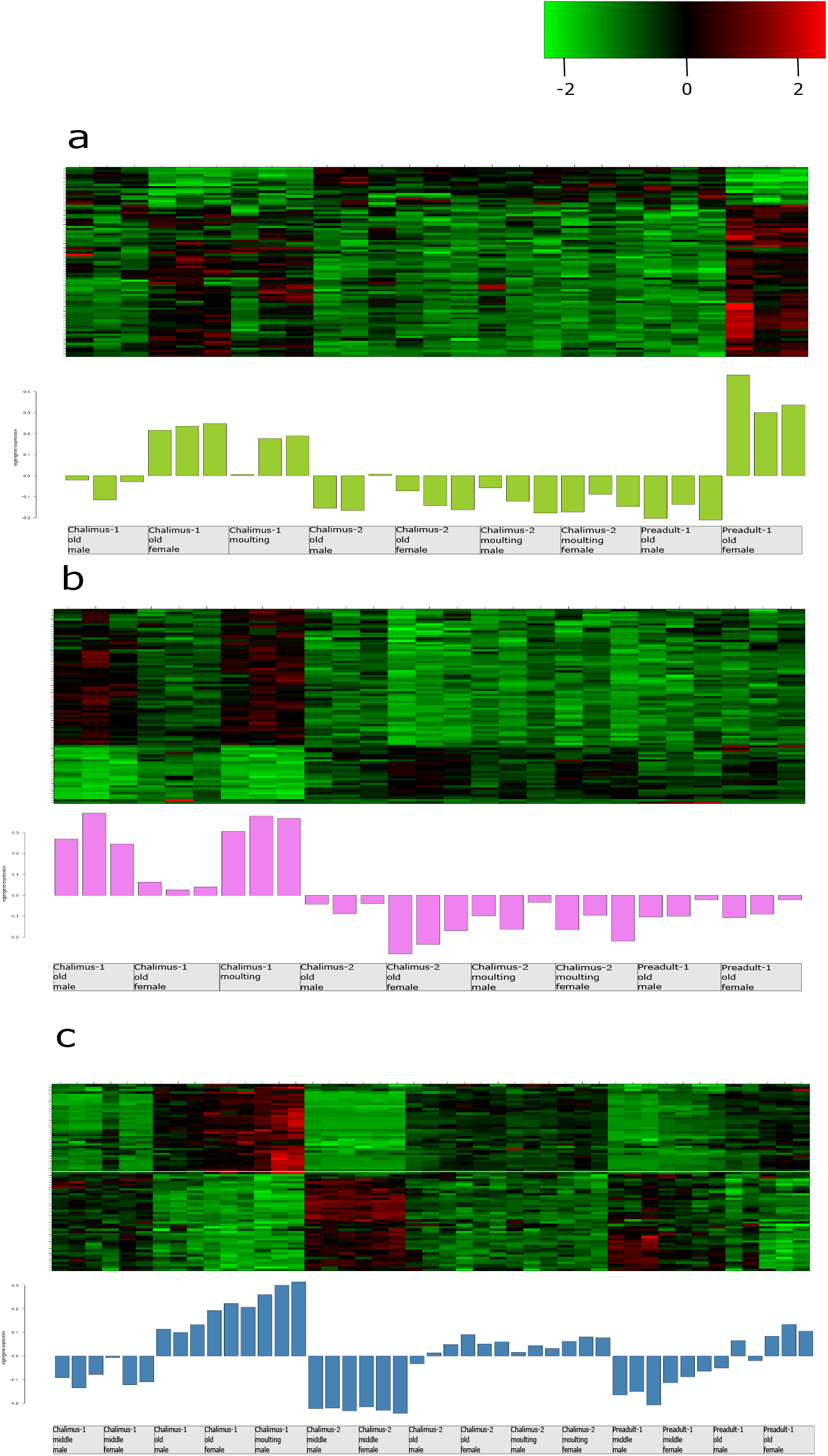
Heatmap of scaled gene expression profiles and barplot of scaled eigengene expression values for module “yellowgreen” extracted from the moulting network (a), module “violet” extracted from the moulting network (b), and module “steelblue” extracted from the global network (c). The heatmap color-codes gene expression values for each gene in a module: higher expression values are represented in red, and lower expression values are represented in greens according to the color legend. The barplot each the heatmap depicts the expression levels of module eigengenes in different samples (x-axis). The barplots are colored based on the name of each module.

Gene expression profiles within a module were strongly correlated. Genes in the module “yellowgreen” tended to be highly expressed in the chalimus-1 old female samples, chalimus-1 moulting samples and preadult-1 old female samples. Genes in the module “violet” were highly expressed in the chalimus-1 male samples and chalimus-1 moulting samples. For the module “steelblue” from the global network, genes were highly expressed in almost all samples from the old and moulting instar stages (except two preadult-1 old male samples), especially in the chalimus-1 moulting samples. The preadult lice with the two genes of the module “yellowgreen” knocked down failed to develop eggstrings. Further study is necessary to understand the role of genes from the module “yellowgreen” in the fecundity of female lice. The topological graph for each of three modules (Figure 6, Additional file 4-6) shows that moulting-associated genes in the module “yellowgreen” and “steelblue” obtained relatively high average ranks, and they were tightly connected with other hubs. In the module “violet”, the moulting-associated gene obtained a low average rank, but the genes annotated as TF obtained high average ranks and were tightly connected with other hubs. For these modules, the proportion of differentially expressed genes was highest in the module “steelblue” identified from the global network.

**Figure 6.**
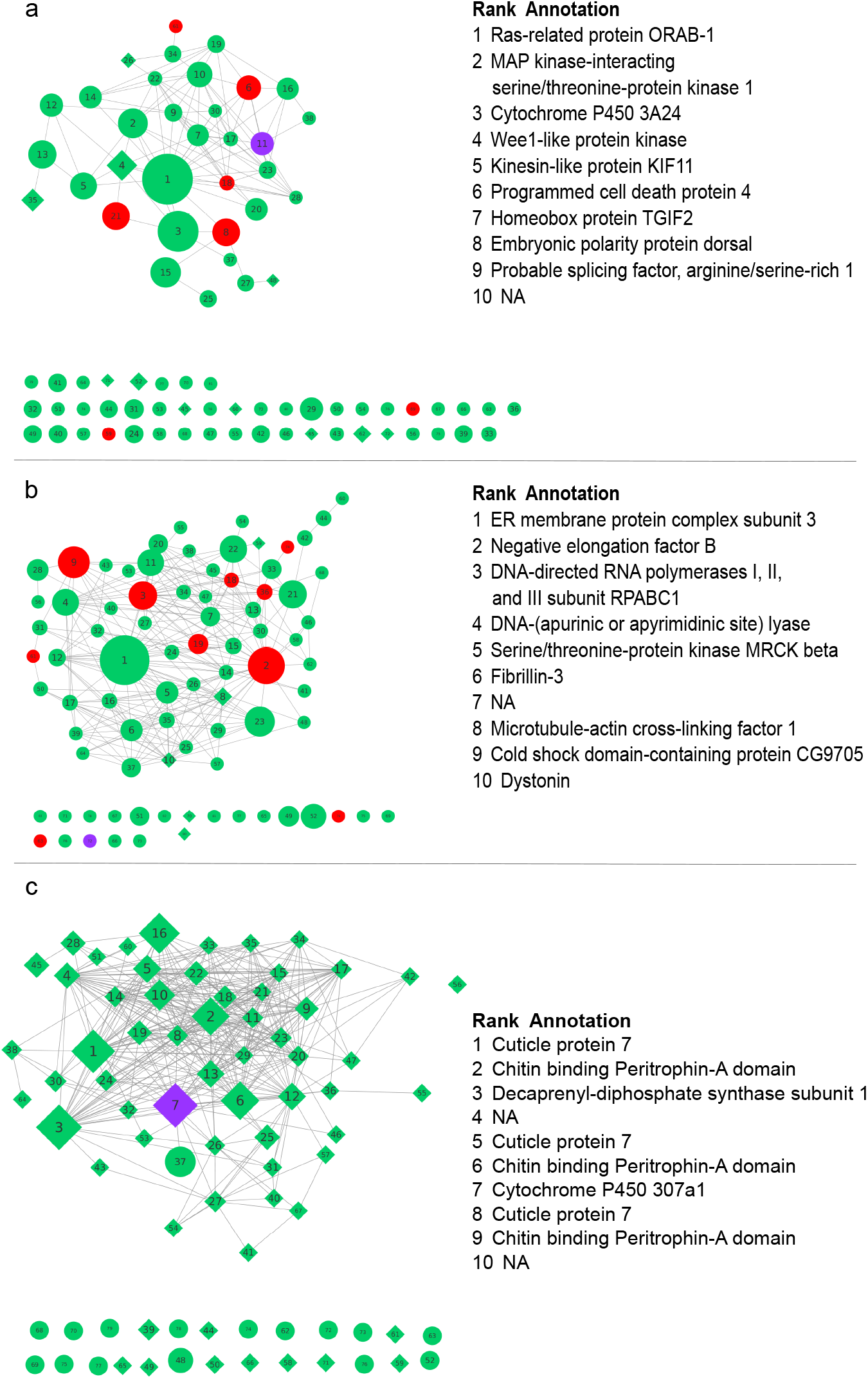
Graph representation of the modules “yellowgreen” extracted from the moulting network (a), “violet” extracted from the moulting network (b), and “steelblue” extracted from the global network (c). Nodes represent genes, edges represent correlation between nodes. For readability, only edges with absolute correlation > 0.84 are drawn. Nodes are labeled based on their average rank over three centrality statistics, node label font size is proportional to average nodes ranks. TF genes are colored in red, purple nodes represent moulting-associated genes from the literature. Differentially expressed genes between non-moulting and moulting groups are displayed as diamond shapes. Node diameter is proportional to weighted betweenness centrality. Annotation information of the top 10 genes based on rank is listed for each module.

The enriched GO terms in the module “yellowgreen” included GO:0008152, GO:0001071 and GO:0005667, indicating nucleic acid transcription factor activity. The enriched GO terms in the module “violet” included GO:0006355, GO:0003677 and GO:0044454, indicating transcriptional regulation. In the module “steelblue”, the most enriched GO terms were GO:0042302, GO:0008061 and GO:0006030, indicating metabolic processes of chitin and cuticle.

## Discussion

RNA interference (RNAi) has enormously facilitated rapid and straightforward analysis of gene function for parasites and other organisms [60–62], and the whole-genome RNAi screen have been successfully applied to detect genes with important functions for many biological processes in C.elegans and mammalian cultured cells [63–65]. Although a robust RNAi method for knocking down salmon louse genes have been established [62], genome-wide RNAi screen for salmon louse is both labour-intensive and time-consuming due to the complex life cycle of salmon louse [3, 4]. Currently, biologists choose RNAi gene candidates subjectively based on their research interests, and little work has yet been carried out to develop bioinformatics methods for objectively predicting salmon louse genes that have a crucial role in biological processes of interest and are likely to show visible phenotypes in RNAi experiments.

In this study, we systematically analyzed the RNA-seq data of salmon lice from different life stages and proposed an approach (a workflow) for identifying important genes involved in the moulting and development of salmon louse (Figure 2). This approach can be used in selecting candidate genes for RNAi experiments. The results of our RNAi experiments and the RNAi records from LiceBase indicate the effectiveness of our approach.

The module preservation analysis allowed us to identify two important genes (EMLSAG00000001458 and EMLSAG00000008959 annotated as RAB1A and DIEXF), and both of the genes were from a non-preserved module (yellowgreen) in the moulting network. The non-preserved modules in the moulting network may be co-regulated and play an indispensable role in moulting or development of the salmon louse. Further studies are required to clarify the biological meaning of the non-preserved modules in the middle network as well as the well-preserved modules between the middle and moulting network.

In the regularized logistic regression analysis, all module eigengenes were calculated using the same method, thus they are on the same scale and it is feasible to identify the most important module by comparing the regression coefficients of eigengenes. We found that the module (steelblue) whose eigengene obtained largest coefficient was enriched for GO categories related to cuticle and chitin metabolic process. All the annotated hubs in this module are associated with chitin binding peritrophin-A domain, cuticle proteins, and cytochrome P450, which have been reported as important proteins for the moulting of arthropods [66–69]. We knocked down a hub (EMLSAG00000004347) with little annotation and observed both reduced fecundity and fragile constituents. Based on RNAi results and the annotations of other hubs in this module, we speculate that gene EMLSAG00000004347 may participate in building of the louse exoskeleton during the moulting process to adult stage. Our approach offers an effective solution in proposing and annotating novel putative genes that play a role in the moulting process of salmon louse. Although we focused on analyzing the modules containing moulting-associated genes due to the limited access to RNAi experiments, the module preservation analysis and regularized logistic regression analysis identify important modules without taking any prior knowledge into account. These methods are suitable to analyze the expression data from less well-annotated organisms.

Instead of focusing on the moulting process directly, the emphasis of homologue-based enrichment analysis is on detecting important modules that are enriched for genes yielding observable phenotypic changes in another species. Four modules were identified by in the first step. Besides the two RNAi experiments performed in this study, RNAi records were found for genes in each of the four modules. Strong RNAi phenotypes were observed on one and three genes in the module “mediumpurple3” and module “turquoise”, respectively. Therefore, homologue-based phenotype enrichment analysis can contribute to rational selection of important modules, especially for studying less well-annotated organisms.

For scale free protein–protein interaction (PPI) networks, many groups have argued that highly connected hub nodes are more likely to be essential than sparsely connected nodes[70–72]. Although the underlying reason is in dispute [73], the centrality-lethality rule [50] has been widely accepted. A recent study on centrality in GCNs arrived at a similar conclusion [74]. Since virtually no PPI data is available for the salmon louse, we focussed on the essentiality of hubs in GCNs instead. Taking the topological characteristics of weighted GCNs into consideration, we used three different methods to identify intramodular hubs. In many cases, the hubs identified with these three measurements were coherent and complementary, enabling us to define absolute hubs. This not only had the advantage of evaluating the intramodular centrality of nodes from different angles, but also increases robustness of our approach. 17 of the 32 moulting-associated genes were detected as intramodular hubs in the three GCNss, and a hexokinase orthologue was found as absolute intramodular hub in the global network and intramodular hub in the other two networks. For the two modules (yellowgreen and violet) from which we chose two hubs in each to knock down, we found that both of the two absolute hubs (EMLSAG00000001458 and EMLSAG00000005299) show stronger phenotypic consequences than the other middle-ranked hub (EMLSAG00000008959 and EML-SAG00000003179). Interestingly, the absolute hub (EMLSAT00000005382) of another module containing moulting-associated transcripts has recently been identified as a novel intestinal heme scavenger receptor with significant phenotypic effect on reproduction [75].

These observations provide support for the significance of intramodular hubs in GCNs. On the other hand, when looking at all other public RNAi experiments in LiceBase [34], we discovered four aditional hub genes that had been tested previously, all of which had highest average rank distributed across four different modules. However, these modules passed none of our criteria for module selection, and negative results had been recorded. We thereby conclude that not all the intramodular hubs may be equally important, even if they have highest ranks, supporting the need for an initial step of module selection. Combining our RNAi results and public records, we argue that our rational approach is more likely to yield genes with measurable phenotypic effect under ablation of gene expression than random selection. Nonetheless, more work is needed to affirm the relationship between centrality and gene essentiality in this organism.

In addition to demonstrating the biological importance of intramodular hubs, RNAi experiments also highlight the role of our selected genes in moulting and development. Ablation of the RAB1A-like gene (EMLSAG00000001458) resulted in reduced survival and fecundity. As a member of the Rab GTPase protein family, Ras-related protein Rab-1A has important roles in many biological processes. Insect Rab proteins have a role in secretion of prothoracicotropic hormone (PTTH) [76, 77], an important regulator of ecdysteroidogenesis [78]. However, few studies have been conducted on Rab proteins in crustaceans. According to our module preservation analysis, another top-scoring hub is annotated as epithelial cell transforming 2 (ECT2). Both, RAB1 and ECT2 are associated with GTPase activity [79–81]. Therefore, the roles of Ras and Rho GTPases in development of the parasite should be further explored.

The top-scoring transcript (EMLSAT00000003849) by RIF analysis is an orthologue of forkhead box protein O (FOXO). The importance of FOXO in metabolism, cellular proliferation, stress tolerance and lifespan has long been recognized [82, 83]. FOXOs are crucial regulators of cellular homeostasis that have a conserved role in modulating organismal aging and fitness [84]. More interestingly, several recent studies have demonstrated that FOXO-like TFs control growth and moulting in insects [85–87]. A homologous TF in *Drosophila melanogaster* (dFOXO) was reported to be involved in regulation of developmental timing through interaction with moulting hormonoe ecdysone [88]. Combining our analysis results with these published papers, we propose that it is worth investigating whether the FOXO-like TF have a crucial role in salmon louse development.

Neither the essential RAB1A-like nor FOXO-like genes, are detected as DE, indicating that DE analysis might not always be the best choice when it comes to identify genes that play a key role in regulating a certain phenotype. In a standard DE analysis, only single genes are taken into account, disregarding possible correlations. On the other hand, some genes linked to a phenotype or disease are not differentially expressed across samples [53, 89], because mutations or post-translational modifications may alter coding potential and function without affecting expression levels [90]. A powerful advantage of network-based analysis is that it can reveal interactions across different groups of samples, even in case of high within-group variability. Furthermore, GCN-based analysis circumvents the multiple testing problem that plagues conventional differential gene expression analysis.

In summary, our results support the hypothesis that GCN-based approaches are effective in identifying genes with association to a phenotype of interest. The widely accepted view that hubs of biological networks are more likely to be essential has for the first time been successfully tested in a marine parasite. Because of the high level of modularity, it was necessary to break down our rational approach of candidate selection by GCN into a two-step process with selecting interesting modules first. In our opinion, improving prioritization of genes is in strong demand parasite functional genomics. This is due to the fact that slow parasite growth as well as labor- and time-intensive handling and collection procedures often render genome-wide functional assays intractable in host-parasite systems. We therefore propose that our selection method may guide gene selection towards candidates with high probability of success in functional studies of sea lice and other parasites. Prospectively, new multi-factorial gene-expression data may also allow to transfer our approach to a broader range of phenotypes.

## Acknowledgements

We would like to thank Heidi Kongshaug, Lars Are Hamre and Per Gunnar Espedal for technical help in the laboratory, and we would like to thank Kjell Petersen for helpful comments and suggestions on the manuscript.

## Funding

This project was funded by the Research Council of Norway, SFI-Sea Lice Research Centre, grant number 203513/ O30. Further, this work was funded by the ELIXIR2 (270068) infrastructure grant from the Research Council of Norway to MD.

### Abbreviations

BC: betweenness centrality
BLAST: Basic Local Alignment Search Tool
BP: biological process
CT: threshold cycle
CC: cellular component
DE: differentially expressed/differential gene expression
dsRNA: double-stranded RNA
DW: differential wiring
fw: forward
FWER: family-wise Type I error rate
GCN: gene co-expression network
GO: gene ontology
KD: knock down
kIM: intramodular connectivity
kME: module eigengene-based connectivity
MF: molecular function
RIF: regulatory impact factor
RNAi: RNA interference
RNA-seq: RNA sequencing
rv: reverse
TF: transcription factor

## Availability of data and materials

The datasets analyzed in this study and code are available from the corresponding author on reasonable request. Supplementary data have been made available in figshare [91]

## Ethics approval and consent to participate

All experiments were performed according to Norwegian animal welfare regulations with the approval of the governmental Norwegian Animal Research Authority (ID7704, no 2010/245410).

## Consent for publication

Not applicable.

## Author’s contributions

ZZ performed bioinformatics analyses and drafted the initial manuscript. MD and CE designed the study, provided and analyzed data. IJ and FN contributed to the design of the study and provided funding. CE performed RNAi experiments and drafted the corresponding methods and results. ZZ, MD and CE edited the manuscript. All authors read, revised and approved the final manuscript.

## Additional Files

Additional file 1 — supplementary methods.pdf

The methods used to estimate the power parameter and calculate the module preservation statistics.

Additional file 2 — analysis results.ods The main analysis results of this study.

Table S1: Homologues-based transcript-phenotype list

Table S2-S3: Analysis results for the modules in the middle network and moulting network, including module size, enriched GO terms with smallest p-values, p-values of enrichment analyses based on homologues, module preservation *Zsummary*, number of known moulting-associated genes, proportion of TF and DE genes.

Table S4: Analysis results for the modules in the global network, including module size, enriched GO terms with

smallest p-values, p-values of enrichment analyses based on homologues, regularized logistic regression coefficients, number of known moulting-associated genes, proportion of TF and DE genes.

Table S5-S7: Module assignment results for the known moulting-associated transcripts in the middle, moulting and global network; the ranks of these transcripts based on three types of centrality measurements within modules; whether they were intramodular hubs or not.

Table S8: Two types of RIF scores for the transcripts annotated as transcription factors

Table S9-S11: Centrality measurements, average ranks and annotations for the nodes in three selected modules (yellowgreen, steelblue and violet).

Table S12: Annotations of the nodes with highest average ranks from the selected modules.

Table S13: Available RNAi results for nodes from the eight selected important modules and nodes with high average ranks from modules without passing any criteria.

Additional file 3 — supplementary figures.pdf

Figure S1: Hierarchical clustering tree (dendrogram) of nodes in the middle, moulting and global network.

Figure S2: Histological sections of adult female louse tissues from control samples and samples with selected genes knock-down.

Figure S3: Histological sections of copepodids sampled three days post infection.

Additional file 4 — yellowgreen.xgmml

Visualization file for the selected important module “yellowgreen” from the moulting network.

Additional file 5 — violet.xgmml

Visualization file for the selected important module “violet” from the moulting network.

Additional file 6 — steelblueGlobal.xgmml

Visualization file for the selected important module “steelblue” from the global network.

Additional file 7 — Adjacency matrices.RData

Adjacency matrices for the middle, moulting and global network.

Supplementary data has been deposited in Figshare DOI: 10.6084/m9.figshare.c.5375315

